# Ecology of the exotic macroalga *Batophora occidentalis* in *Posidonia oceanica* meadows and other native benthic habitats

**DOI:** 10.1101/2025.11.20.689540

**Authors:** Andrea Anton, Silvia Paoletti, Lidia Cucala-Garcia, Carlos Morell, Sara Muñiz-Quintana

## Abstract

Marine invasive macroalgae can be particularly detrimental to native ecosystems. The exotic green macroalga *B. occidentalis* was first reported in 2020 in Estany des Peix lagoon in Formentera (Spain). In 2023 and 2024, our surveys revealed that *B. occidentalis* had spread to 100% of the surveyed locations within the lagoon. In 2024, *B. occidentalis* reached an average cover of 30% in *Cymodocea nodosa* meadows and 26% in *Caulerpa prolifera* meadows, with respective biomasses of 52 g m^−1^ and 44 g m^−1^, making it the most abundant macrophyte in these two native benthic habitats. We also found exotic *B. occidentalis* growing epiphytically on the leaves and rhizomes of *Posidonia oceanica* within the lagoon, with an average cover of 12% in 2023 and 16% in 2024, and a biomass of 3 g m^−1^ in 2024. Notably, it covered nearly 25% of the *P. oceanica* leaf surface and reached up to five times the weight of individual leaves. Alarmingly, we report the presence of a *B. occidentalis* specimen on a *P. oceanica* shoot in a nearby meadow outside the lagoon, signaling a potential spread beyond its current range. Additionally, *B. occidentalis* colonized various natural and hard substrates—including boat hulls, as it was found growing on 17 % of boats anchored in the only marina in the lagoon. With this study, we aim to encourage prompt action from local and regional governments. We may be witnessing the early stages of a broader *B. occidentalis* expansion, highlighting a critical moment for implementing early management measures such as monitoring, and, if possible, containment within the lagoon.

## INTRODUCTION

The introduction of species is an increasingly significant environmental challenge, with an estimated 35% rise in the number of emerging exotic species globally by 2050 (Seebens 2020; Anton 2021). The Mediterranean Sea, home to more than 1,000 reported exotic species (Zenetos et al. 2022), is the most invaded marine region in the world (Roy et al. 2024), where marine exotics can have a significant ecological impact on biodiversity and endemic species (Marbà et al. 2014; Geraldi et al. 2020; Puentes and Anton 2025; Pizarro-Borrull et al. 2025). This high number of introductions is driven by various factors, including the opening of the Suez Canal in 1869, increased shipping activity, and accidental introductions from aquaculture facilities (Roy et al. 2024). Among these, marine exotic macrophytes are responsible for the largest environmental impacts on native species (Anton et al. 2019), primarily by negatively affecting other (native) primary producers (Anton et al. 2020a). For example, the invasive macroalgae *Lophocladia lallemandii* has been shown to increase the mortality of seagrass *Posidonia oceanica* shoots by 2.5 to 5 times (Marbà et al. 2014). The Mediterranean Sea is home to an estimated 118 introduced macrophyte species, including macroalgae and seagrasses, with an average of 22 new introductions per decade since 1990 (Wesselmann et al. 2024). One such recent introduction is the green macroalga *Batophora occidentalis* J. Agardh, 1854 (Chlorophyta: Dasycladales).

*Batophora occidentalis* was first detected in the Mediterranean in the spring of 2020, in the Estany des Peix in the island of Formentera (Balearic Islands; Spain) (Ballesteros 2020), where its distribution was afterwards updated in 2023 (Forteza et al 2024). The native range of the genus *Batophora* is the tropical and subtropical western Atlantic (Ballesteros 2020). Outside this native range, it was first reported in the Canary Islands, Spain, between 1990-1992 (Afonso Carrillo et al. 1993) and in the Chesapeake Bay, USA, in 2015 (Hall and Schneider 2023). Another Mediterranean record of the genus comes from the Mar Menor lagoon in the Murcia region in southern Spain (Terradas-Fernández et al. 2022). The two Mediterranean detections of *B. occidentalis* occurred almost simultaneously: in Formentera in May 2020 and Mar Menor in November 2021, with specimens from both introductions having overlapping morphological traits (Ballesteros 2020; Terradas-Fernández et al. 2022). It has been hypothesized that both populations may be linked to the same introduction event (Terradas-Fernández et al. 2022), with a likely pathway of introduction to Formentera being shipping, particularly luxury yachts that overwinter in the Caribbean and travel to the Mediterranean in the summer (Ballesteros 2020).

There are three species or varieties within the genus *Batophora*: *Batophora oerstedii, Batophora occidentalis* and *Batophora occidentalis* var. *largoensis* (Rodriguez-Reyes et al. 2018). These species have been taxonomically recognized based on morphological characteristics such as the size of the algae, the location of the whorls along the main axis, the position and size of the gametophores, and the size and shape of the gametangia (Rodriguez-Reyes et al. 2018). However, there is no definitive scientific consensus on how these traits define the species, which complicates their identification. Terradas-Fernández et al. (2022) identified the *Batophora* species in Mar Menor as either *Batophora occidentalis* or *Batophora occidentalis* var. *largoensis* based on morphological data. Similarly, Hall and Schneider (2023) used morphological traits to identify the introduction of *Batophora oerstedii* in the Chesapeake Bay. *Batophora* from Estany des Peix has also been identified tentatively as *Batophora occidentalis* var. *largoensis* based on morphological characteristics, though the author noted that the identification was inconclusive (Ballesteros 2020). Lombardo et al. (2025) refer to the species of *Batophora* in Formentera as *B. occidentalis* and we also do so in this study.

Here, we describe the extent and ecological characteristics of the *B. occidentalis* invasion in Estany des Peix in Formentera 3 to 4 years after its initial report (Ballesteros 2020). Specifically, we: 1) assessed the presence of *B. occidentalis* around the perimeter (and nearby locations outside) of the lagoon during the late summers of 2023 and 2024, 2) documented, with photographs, the presence of *B. occidentalis* on various habitats, including natural substrates, artificial structures, and boats, 3) quantified using video-tapes the *B. occidentalis* benthic cover in two locations (a *Posidonia oceanica* meadow and sandy bottom) inside the lagoon in 2023, and in four locations in 2024 (a meadow of *Cymodocea nodosa*, a meadow of *Caulerpa prolifera*, and two *P. oceanica* meadows located inside and outside the lagoon, respectively), 4) measured the biomass of *B. occidentalis* growing on the leaves of these three macrophyte species and/or in the surrounding sediment within the meadows in 2024, and 5) quantified the presence of *B. occidentalis* attached to the hulls of the boats anchored in the main marina of the lagoon.

## METHODS

To assess the distribution of *B. occidentalis* around the perimeter of the lagoon Estany des Peix (38.72618, 1.41200; Formentera Island, Spain; Figure 1), we conducted visual surveys on September 27^th^ and 28^th^ 2023 and 2^nd^ - 3^rd^ of October 2024. A total of 12 and 44 study locations were surveyed in 2023 and 2024, respectively. In 2023, eight locations were surveyed inside the lagoon (locations 1-8; Figure 1A), one location at the entrance (location 9; Figure 1A), and three locations outside the lagoon (locations 10-12; Figure 1A). Inside the lagoon, seven locations were situated around the perimeter at depths of less than 1 meter, whereas one location (location 3; Figure 1A) was surveyed at approximately 2 meters depth. In 2024, 34 locations were surveyed around the perimeter of the lagoon and 10 were surveyed outside the lagoon, all at less than 1.5 meters depth (locations 13-55; Figure 1B). At each location we visually inspected approximately a 50-meter stretch to determine the presence of *B. occidentalis*. In 2023 we measured temperature in the two study locations (locations 1, and 2; Figure 1A) while in 2024 we measured temperature, salinity, and dissolved oxygen (mg l^-1^ and %) in four study locations (locations 13-16; Figure 1B) using a ProSolo (YSI Xylem) handheld probe. In 2024, we quantified the number of boats tied to the dock in the marina at the Estany des Peix that had *B. occidentalis* growing on their hulls by visually inspecting all the boats from the dock.

**FIGURE 1.**
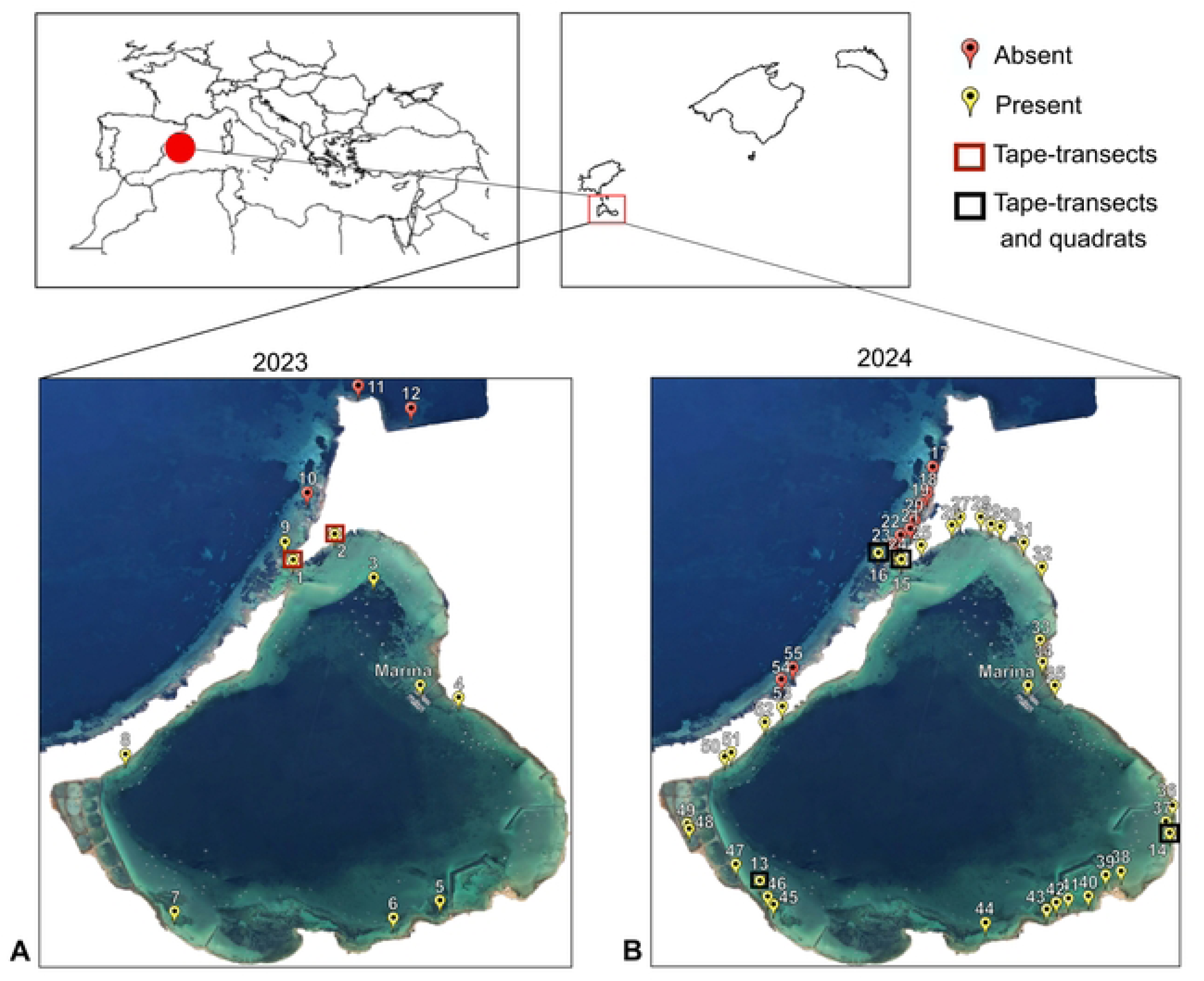
Map of the study site, the Estany des Peix lagoon, in Formentera Island (Balearic Islands, Spain) in **A**) 2023 and **B**) 2024. Points (1-55) indicate the presence (green) or absence (red) of *B. occidentalis* assessed visually in eight locations inside the lagoon and four locations outside the lagoon in 2023, and 34 locations inside the lagoon and 10 locations outside the lagoon in 2024, where pictures were also taken. In 2023, locations 1 and 2 (red squares) were recorded with tape-transects for % benthic cover. In 2024, locations 13-16 (blue squares) were recorded with tape-transects for % benthic cover and sampled with quadrats for biomass analysis.

In 2023, we quantified *B. occidentalis* and main macrophyte % benthic cover at two study locations (location 1 and 2, Figure 1A) corresponding to a sandy bottom habitat and a *P. oceanica* meadow by deploying six and two 10-meter-long transect tapes, respectively, and conducting video recordings along the tape using a GoPro Hero 7 camera. Similarly in 2024, we quantified benthic cover at four study locations (location 13-16, Figure 1B) corresponding to three benthic habitats inside the lagoon (meadows of *P. oceanica*, *C. nodosa* and *C. prolifera*) and a benthic habitat outside the lagoon (a meadow of *P. oceanica*) by deploying and recording three 10-meter-long transect tapes at each location. For each transect, the timestamp was noted, and a video of 98 ± 17 seconds was taken while hovering over the transect at a close distance. In the laboratory, coverage annotations were made using a point transect method at every 10 cm line mark (n=100 per transect; following Anton et al. 2014, 2020b). The seafloor was classified in one the following categories: (1) meadow (either *C. prolifera*, *C. nodosa* or *P. oceanica*, depending on the biogenic habitat being assessed), (2) *B. occidentalis* on sediments, (3) *B. occidentalis* on alive leaves, (4) dead matte, (5) rock or (6) sand. Based on the knowledge acquired on the species occurrence from the analysis of the quadrat samples (see below), *B. occidentalis* was either classified as (2) *B. occidentalis* on sediments when it was independently hooked on the substrate, even if in close proximity to another species, or (3) *B. occidentalis* on leaves when it was attached to the seagrass/seaweed structures. When the coverage was classified as (3) *B. occidentalis* on leaves, despite it also implied coverage by the specific meadow type below, the point was only annotated once to be able to reach 100 % coverage. Dead matte represented the substrate made of remaining underlying *P. oceanica* rhizomes, roots, and sediment particles when the above canopy has died. Both sides of the 10 cm mark line were considered to aid classification, however in case of different underlying coverage, the left side of the mark line was consistently chosen. Whenever the underlying cover changed depending on frame perspective or the image was of lesser quality due to movements and glare, annotations were made when the mark line was on focus and/or placed at the centre of the frame. Coverage percentages were calculated as follow:

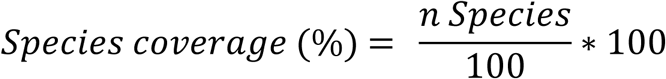

Total *B. occidentalis* coverage was calculated as the sum of *B. occidentalis* on sediments and *B. occidentalis* on leaves. Total meadow coverage was calculated as the sum of meadow and *B. occidentalis* on leaves as the latter implied an underlying meadow coverage.

In 2024, three quadrat samples of 10 x 10 cm were collected at the 1, 5, and 9 m points on each transect of the four study locations (totalling 36 quadrat samples). Quadrat samples were collected into zip lock bags and conserved at −18°C until processed. For each sample, meadow density was estimated from the number of shoots, except for the *C. prolifera* habitat where total number of leaves were noted. Specimens were sorted, and wet and dry weights were measured separately for 1) unimpacted meadow biomass, 2) impacted meadow biomass (i.e., with *B. occidentalis* on leaves), 3) *B. occidentalis* biomass independently hooked on the sediments, and 4) biomass of *B. occidentalis* on leaves. Wet weight was measured after draining on absorbent paper using a precision balance (resolution 100 µg). Dry weight was measured after drying at 60°C for three days. The total number of *B. occidentalis* stalks and the proportion of fertile ones were noted for both above categories. For the *C. nodosa* and *P. oceanica* habitats, the number of leaves per shoot was also noted. For the *P. oceanica* habitat, shoots were processed individually. Position in the shoot (i.e., chronologically from the centre), total length, and minimum height of the *B. occidentalis* attachment were noted for each invaded leaf. The way rhizoids attached on the leaf (i.e., directly on leaf tissue, or on epiphytes) was noted. Illustrative specimens were photographed using an Olympus Tough TG-5 camera, and observed through a stereomicroscope and a ZEISS Axio Zoom V16 digital microscope. All statistical analysis were done using R. The differences in mean coverage among biogenic habitats were compared using a one-way ANOVA test followed by a post-hoc Tukey HSD test. When ANOVA assumptions (Kozak and Piepho, 2017) were not met, non-parametric Kruskal-Wallis test followed by Dunn’s test were used instead. The differences in biomass among benthic habitats were compared using a linear mixed model (*lme()* function using package *nlme*) with transect ID as random effect, followed by a post-hoc Tukey HSD (*emmeans()* function using package *emmeans*). Residual plots were used to check for violations of normality on residuals (Schielzeth et al., 2020). The relationships between continuous variables were explored using linear regression modelling (*lm()* function using package *stats*).

## RESULTS

The average temperature in the lagoon during the surveys in 2023 and 2024 was 26.5 and 25.1 ^°^C, respectively, while salinity and dissolved oxygen were on average 38.9 ppt and 7.6 mg l^-1^ (or 116 % saturation) in 2024, respectively. Our findings revealed that *B. occidentalis* was present in all the locations surveyed along the perimeter of the lagoon in 2023 and 2024 at <1 m depth (Figure 1A and B) and in the additional surveyed location within the lagoon in 2023 at approximately 2 m depth (location 3; Figure 1A). We also observed *B. occidentalis* at the entrance of the lagoon in rock pools in 2023 (location 9; Figure 2A) but not in the visual surveys at the entrance of the lagoon in 2024 location 16; Figure 1B). *Batophora occidentalis* was visually not present on the other three and 10 locations outside of the lagoon in 2023 and 2024, respectively (locations 10-12, 16, 18-23, 54, and 55, Figure 1A and B).

**FIGURE 2.**
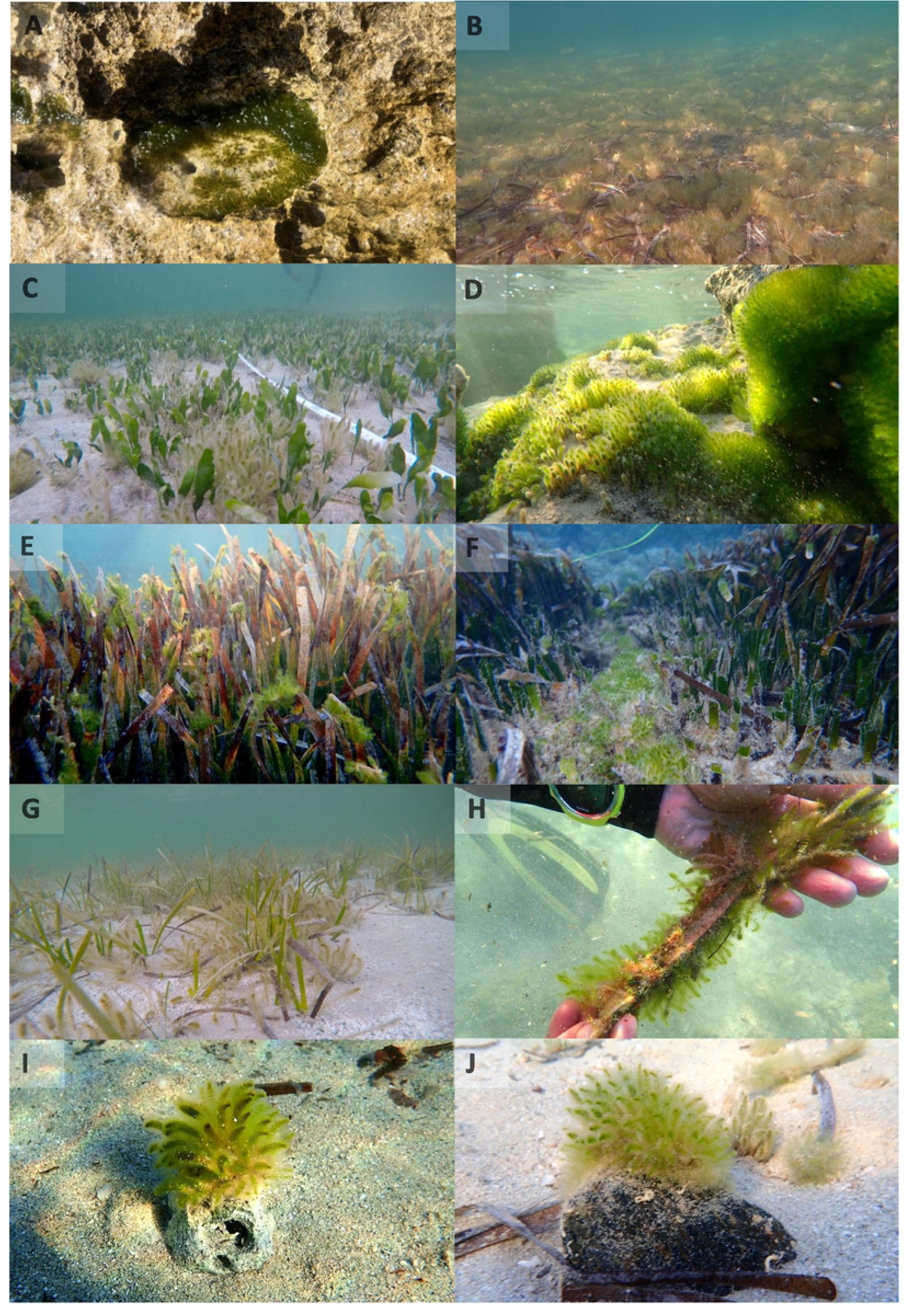
Exotic macroalgae *B. occidentalis* (A) growing on rock pools outside the lagoon, (B) forming extensive meadows growing mainly on *Posidonia oceanica* detrital leaves (2 m depth approx.), (C) growing on meadows of the macroalgae *Caulerpa prolifera*, (D) proliferating on pebbles and rocky outcrops, (E) growing on the leaves of the seagrass *Posidonia oceanica*, (F) expanding on the rhizome of sparse areas within *Posidonia oceanica* meadows, (G) proliferating on meadows of the seagrass *Cymodocea nodosa*, (H) growing on detrital leaves of *Posidonia oeanica*, and (G-H) growing on natural hard substrates in sandy areas (e.g., old sea shells; G and H) in the Estany des Peix in the island of Formentera in September 2023 and October 2024. Photo credit: Andrea Anton.

*B. occidentalis* grows, as documented on pictures, on various natural habitats (Figure 2) including the leaves and rhizomes of healthy seagrass *P. oceanica* meadows (Figure 2E and F), detrital leaves of *P. oceanica* (Figure 2H), in the sediment of seagrass *C. nodosa* meadows (location 5; Figure 2G), rocks and rubble (Figure 2D), and other biogenic hard structures scattered in the sandy bottom such as seashells (Figure 2I, and J). *Batophora occidentalis* was also commonly found on man-made structures including wooden harbours (Figure 3A), marine debris (e.g., cans, plastic pieces; Figure 3B), boat anchor chains (Figure 3C), and boat hulls (Figure 3D). We found *B. occidentalis* growing on 17.4 % (11 out of 63) of the hulls of the boats anchored to the main marina of the lagoon (38.727519, 1.415702). Additionally, we found clumps of *B. occidentalis* floating throughout the lagoon.

**FIGURE 3.**
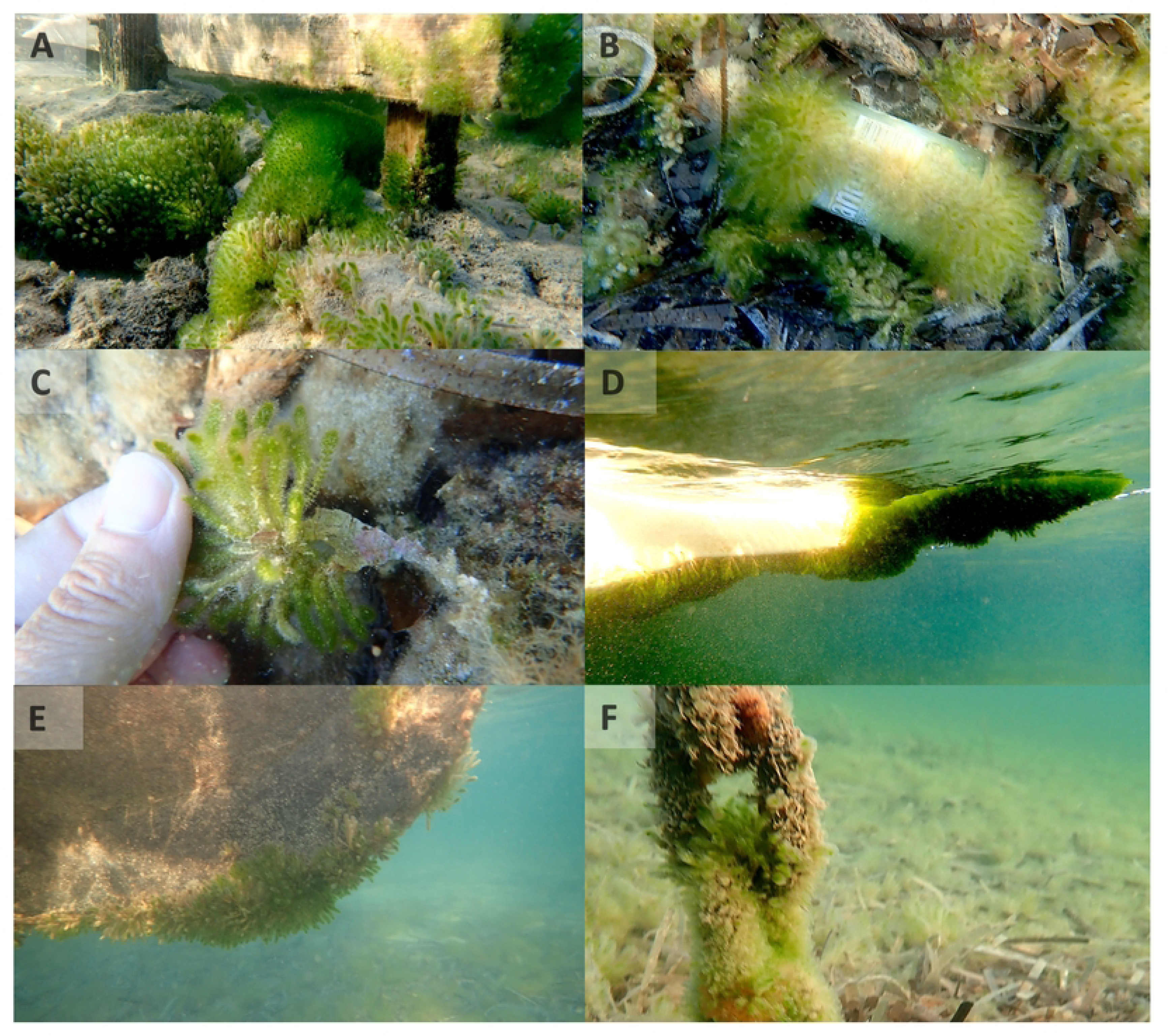
Exotic macroalgae *B. occidentalis* growing on man-made structures such as wooden harbours (A), marine debris (e.g., soda can; B), boat anchor chains (C) and boat hulls (D) in the Estany des Peix in the island of Formentera in September 2023. Photo credit: Andrea Anton.

The tape-transects recorded in 2023 highlighted a widespread distribution of *B. occidentalis* in both sand and a *P. oceanica* meadows inside the lagoon, with 15.8 ± 7.3 % and 17.5 ± 14.8 % of average total coverage, respectively (Figure 5A, Figure S2 and Table 1). In sand (62.8 ± 15.0 % sand coverage; location 1; Figure 1A and Table 1), *B. occidentalis* was observed mainly attached to detached debris and detrital *P. oceanica* leaves. In the *P. oceanica* meadow inside the lagoon (location 1; Figure 1A), the percentage of leaves containing *B. occidentalis* out of the total meadow coverage in 2023 was similar to that of 2024, with 25.0 ± 8.9 % and 22.9 ± 3.0 % of leaves covered with *B. occidentalis*, respectively (Table 1).

**TABLE 1.** Tape-transects mean ± SD coverage percentage (% summing to 100) in 2023 and 2024 of native seagrass/seaweed species, exotic *Batophora occidentalis* on sediments or on leaves, and remaining sediment types for each benthic habitat (n=12).

| Habitat type | Year | Location | Native macrophyte | *Batophora* on sediments | *Batophora* on leaves | *Posidonia* matte | Rock | Sand |
| --- | --- | --- | --- | --- | --- | --- | --- | --- |
| Sand bottom | 2023 | Lagoon | 0.0 ± 0.0 | 15.8 ± 7.3 | 0.0 ± 0.0 | 21.3 ± 9.2 | 0.0 ± 0.0 | 62.8 ± 15.0 |
| *Posidonia oceanica* | 2023 | Lagoon | 29.5 ± 23.3 | 5.5 ± 2.1 | 12.0 ± 12.7 | 3.5 ± 4.9 | 32.5 ± 19.1 | 17.0 ± 14.1 |
| *Caulerpa prolifera* | 2024 | Lagoon | 18.0 ± 11.5 | 25.7 ± 4.7 | 0.0 ± 0.0 | 0.0 ± 0.0 | 2.3 ± 2.1 | 54.0 ± 8.2 |
| *Cymodocea nodosa* | 2024 | Lagoon | 15.3 ± 6.5 | 30.0 ± 3.6 | 0.0 ± 0.0 | 0.0 ± 0.0 | 0.0 ± 0.0 | 54.7 ± 9.3 |
| *Posidonia oceanica* | 2024 | Lagoon | 54.7 ± 3.2 | 0.7 ± 0.6 | 16.3 ± 3.2 | 8.0 ± 1.7 | 13.7 ± 2.5 | 6.7 ± 1.2 |
| *Posidonia oceanica* | 2024 | Open sea | 81.7 ± 6.8 | 0.0 ± 0.0 | 0.0 ± 0.0 | 10.6 ± 6.5 | 1.3 ± 1.5 | 6.3 ± 3.8 |

In 2024, tape-transects indicated that the total meadow coverage across the sampled benthic habitats ranged between 23 and 82 %, with *C. prolifera* covering 23.0 ± 12.5 % of the seafloor, *C. nodosa* 25.0 ± 12.0 %, *P. oceanica* inside the lagoon 71.0 ± 5.6 %, and *P. oceanica* in the open sea 81.7 ± 6.8 % (Figure 4B and Table 1). *Batophora occidentalis* was present in three out of four locations (Figure 4C), as it was not observed in the open sea *P. oceanica* meadows outside of the lagoon. *Batophora occidentalis* coverage was 25.7 ± 4.7 % in *C. prolifera* habitats, 30.0 ± 3.6 % in *C. nodosa habitats*, and 17.0 ± 2.7 % in *P. oceanica* habitats inside the lagoon; in the first two cases higher on average than the native habitat-forming species (Figure 4B and Table 1). Post-hoc pairwise comparisons using Tukey HSD test indicated that total *B. occidentalis* coverage were significantly different among habitats, except between *C. prolifera* and *C. nodosa* (Figure 4C). *Batophora occidentalis* was only observed on leaves in *P. oceanica*, in coverage % significantly different from all other habitats (*p* < 0.0005, one-way ANOVA; Table 1, Figure 4D).

**FIGURE 4.**
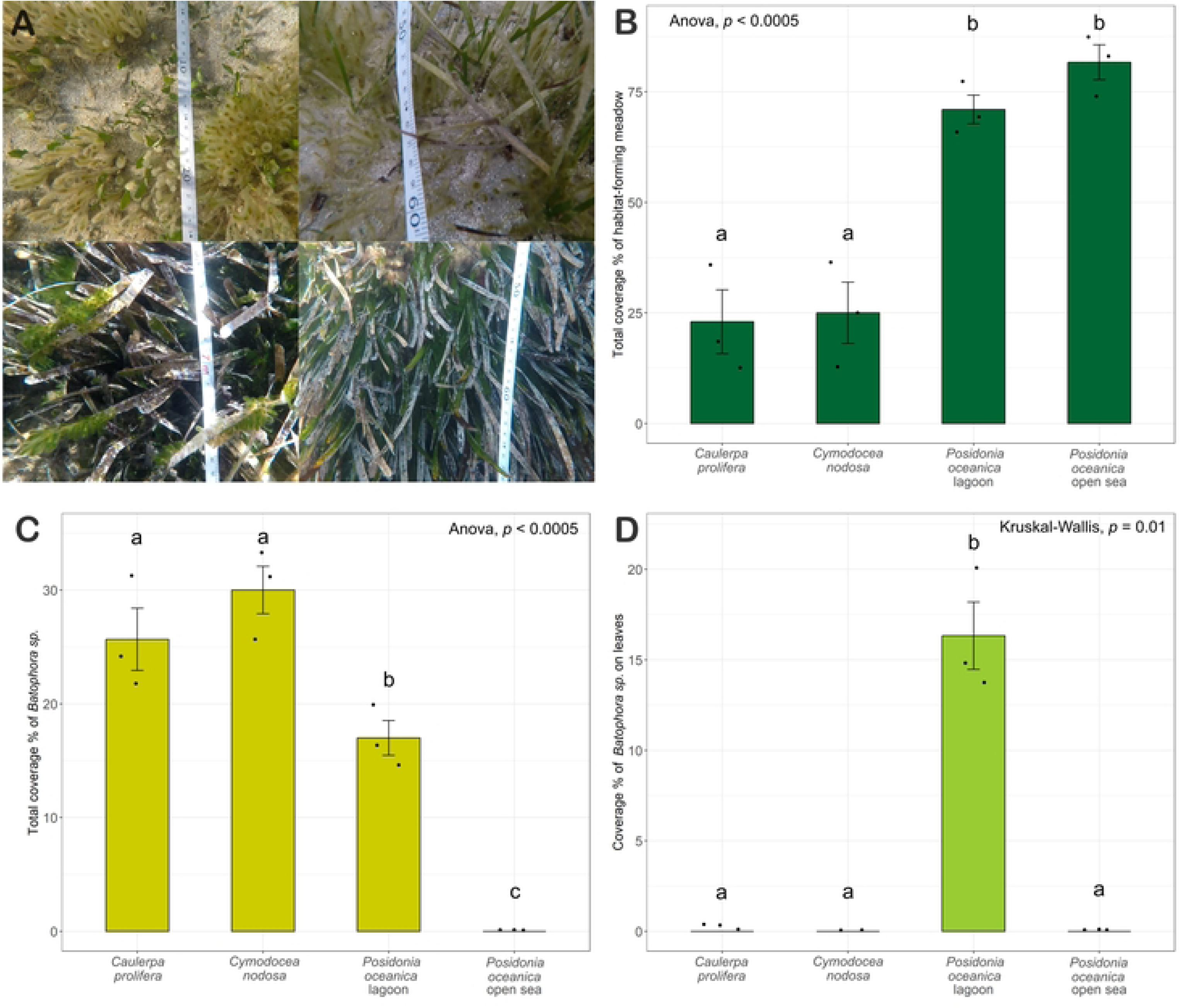
*Batophora occidentalis* and main macrophyte % benthic coverage from tape-transects in 2024. **A)** Examples of each benthic habitat sampled by video recordings; **top left**: *Caulerpa prolifera* habitat (location 13 on Figure 1B), **top right**: *Cymodocea nodosa* habitat (location 14 on Figure 1B), **bottom left**: *Posidonia oceanica* habitat inside the lagoon (location 15 on Figure 1B), and **bottom right**: *Posidonia oceanica* habitat outside the lagoon in the open sea (location 16 on Figure 1B). Barplots of (**B**) total meadow coverage (%), (**C**) total *Batophora* coverage (%), and (**D**) coverage (%) of *B. occidentalis* on leaves, in each benthic habitat in 2024. The results of the one-way ANOVA and of Kruskal-Wallis test are shown, while letters indicate statistically significant differences in the means tested by post-hoc pairwise comparison using Tukey HSD and Dunn’s test, respectively.

In 2024, *B. occidentalis* was found in all quadrat samples collected inside the lagoon (location 12-15; Figure 1B) except for one sample in the *C. nodosa* habitats. One *B. occidentalis* stalk was found on one leaf of *P. oceanica* outside the lagoon in the open sea (location 16; Figure 1B and Figure S3). The total biomass of *B. occidentalis* regardless of where it was attached significantly differed between *C. nodosa* and *P. oceanica* habitat in the open sea (*p* = 0.03, Tukey’s test). The biomass of *P. oceanica* both inside the lagoon and in the open sea were significantly higher than the biomass of *C. prolifera* and *C. nodosa* inside the lagoon (Tukey’s test, Figure 5A). In the locations with *C. prolifera* and *C. nodosa* meadows, the near totality of *Batophora* was found attached independently to the sediments, achieving higher biomass than the native habitat-forming species (Table 2, Figure 5B). The size of *B. occidentalis* bundles varied across samples (Table 2); the biggest ones by weight and number of stalks were observed in *C. nodosa* habitat (Figure 5C), however the bundles in *C. prolifera* habitat had a higher ratio of fertile stalks (23.2 ± 15.6 %) compared to *C. nodosa* and *P. oceanica* habitats (15.9 ± 21.4 % and 2.5 ± 5.3 %, respectively). In *P. oceanica* habitat inside the lagoon, the near totality of *B. occidentalis* was found attached on *P. oceanica* leaves. The biomass of *B. occidentalis* on leaves ranged between zero and five times the weight of the colonized leaf (0 - 5.2 g g^−1^ DW), with an average of 0.50 ± 0.96 g g^−1^ DW (Figure 5F). *Batophora occidentalis* was observed attached to the leaves of species other than *P. oceanica* in only four instances: once on *C. nodosa*, twice on *C. prolifera*, and once on a single *Halimeda tuna* specimen found while sampling the *C. prolifera* location. As such instances were rare, only photographic evidence is reported in the Supplementary materials (Figure S4).

**FIGURE 5.**
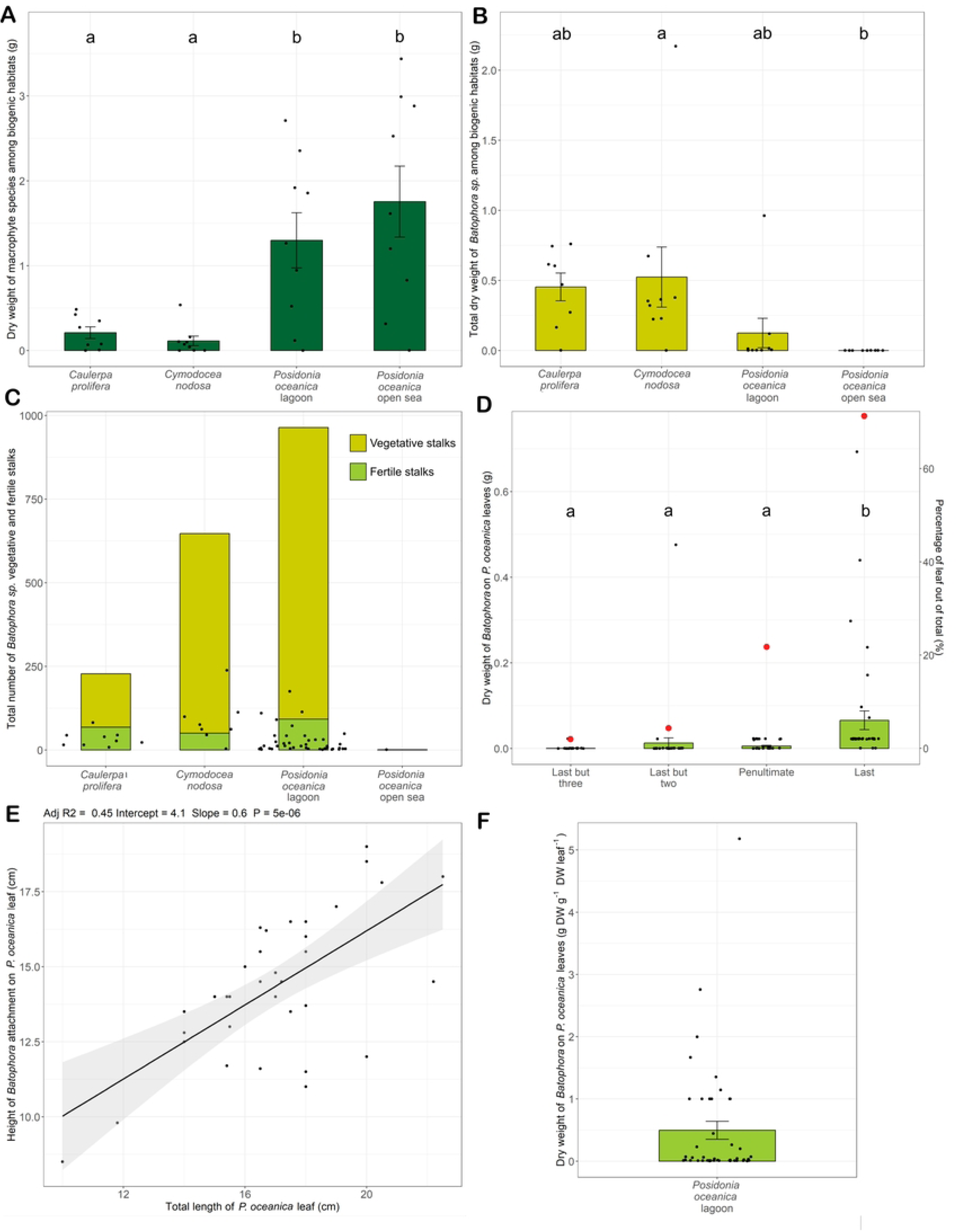
Differences in mean meadow biomass (**A**), mean total *B. occidentalis* (**B**), and sum of *B. occidentalis* vegetative and fertile (with gametophores) stalks among biogenic habitats in 2024 (**C**). **D**) Distribution of *B. occidentalis* biomass on *P. oceanica* leaves inside the lagoon as a function of leaf position in the shoot, with red points indicating the proportion (%) of leaves with *B. occidentalis* for each position among the total number of shoots covered with *B. occidentalis*. **E**) Relationships between the height of *B. occidentalis* attachment on the leaf and the total length of the latter. **F**) Epiphytic load of *B. occidentalis* on leaves expressed in g DW g^-1^ DW leaf. Barplots show the results of the linear mixed models with letters indicating statistically significant differences in the means, while the scatterplot shows the results of the linear regression model.

**TABLE 2.** Summary of data (mean ± SD) obtained from the quadrant samples analysis in 2024. Meadow density is presented as n° shoots/0.01 m^2^, except for *C. prolifera* habitat which is expressed in n° leaves/0.01 m^2^. Biomass is presented as dry weight in g/0.01 m^2^. *Batophora occidentalis* biomass is presented separately as biomass independently hooked on the sediments (per sample) and biomass on leaves (per leaf), respectively. The sample size available for computing each average value is indicated below.

| Habitat type | Location | Meadow density | Meadow biomass | N° leaves per shoot | Shoot biomass | *Batophora* on sediments biomass | *Batophora* on sediments n° stalks (fertile) | *Batophora* on leaves biomass | *Batophora* on leaves n° stalks (fertile) | *Batophora* biomass on covered leaves |
| --- | --- | --- | --- | --- | --- | --- | --- | --- | --- | --- |
| *Caulerpa prolifera* | Lagoon | 17.38 ± 15.29<br>(n = 8) | 0.21 ± 0.20<br>(n = 8) | - | - | 0.44 ± 0.28<br>(n = 8) | 42 ± 23<br>(10 ± 6)<br>(n = 8) | 0.04 ± 0.05<br>(n = 2) | 15 ± 7<br>(3 ± 4)<br>(n = 2) | 0.04 ± 0.05<br>(n = 2) |
| *Cymodocea nodosa* | Lagoon | 3.56 ± 3.68<br>(n = 9) | 0.11 ± 0.17<br>(n = 9) | 4.04 ± 0.98<br>(n = 23) | 0.12 ± 0.13<br>(n = 7) | 0.52 ± 0.64<br>(n = 9) | 99 ± 65<br>(7 ± 5)<br>(n = 8) | < 0.0005<br>(n = 1) | 3<br>(2)<br>(n = 1) | < 0.0005<br>(n = 1) |
| *Posidonia oceanica* | Lagoon | 10.12 ± 5.77<br>(n = 9) | 1.30 ± 0.98<br>(n = 9) | 6.05 ± 2.13<br>(n = 78) | 0.13 ± 0.09<br>(n = 78) | 0.002 ± 0.007<br>(n = 9) | 49<br>(3)<br>(n = 1) | 0.03 ± 0.08<br>(n = 44) | 25 ± 37<br>(2 ± 7)<br>(n = 44) | 0.06 ± 0.09<br>(n = 44) |
| *Posidonia oceanica* | Open sea | 11.33 ± 6.52<br>(n = 9) | 1.75 ± 0.25<br>(n = 9) | 6.00 ± 1.85<br>(n = 102) | 0.15 ± 0.13<br>(n = 102) | 0.00 ± 0.00<br>(n = 9) | - | < 0.0005<br>(n = 1) | 1<br>(0)<br>(n = 1) | 0.02<br>(n = 1) |

In the *P. oceanica* meadows inside the lagoon, *B. occidentalis* was found to be mainly attached to the last (71 % of cases) and penultimate (22 % of cases) leaves of the shoots (Figure 5D), with a significantly higher biomass on the oldest leaf (*p* = 0.0002, Tukey’s test) than the subsequent younger leaves. However, it was often the case that when *B. occidentalis* was found on the penultimate leaf, the oldest leaf was broken (and short), and as such, they could have hosted *B. occidentalis* previously on their missing tip. *Batophora occidentalis* was always found at the tip of the leaf; with the height of the attachment resulting significantly correlated with the length of the leaf (Adj-R^2^ = 0.45, *p* < 0.0005, linear regression model; Figure 5E).

A look at the leaves of *P. oceanica* covered with *B. occidentalis* revealed that the tops of the leaves were covered by a dense mesh of *B. occidentalis* brown rhizoids from which the vegetative (free of spherical gametophores) and fertile (with spherical gametophores) stalks come out (Figure 6). The rhizoid system and the stalk bases were observed to attach to the epiphytes growing on the leaf. Due to the fluffy nature of the stalks, sediment was observed in the bundles. In the lower parts of the leaves, fewer and smaller stalks were observed, although the rhizoid system still appeared to be quite extensive wherever epiphytes were present (Figure A to D). The bottom panels of Figure 6 (E-H) highlighted how an array of spot-like green patches not visible to the naked eye seem to be the precursor of new rhizoids and stalks in the lower parts of the leaf.

**FIGURE 6.**
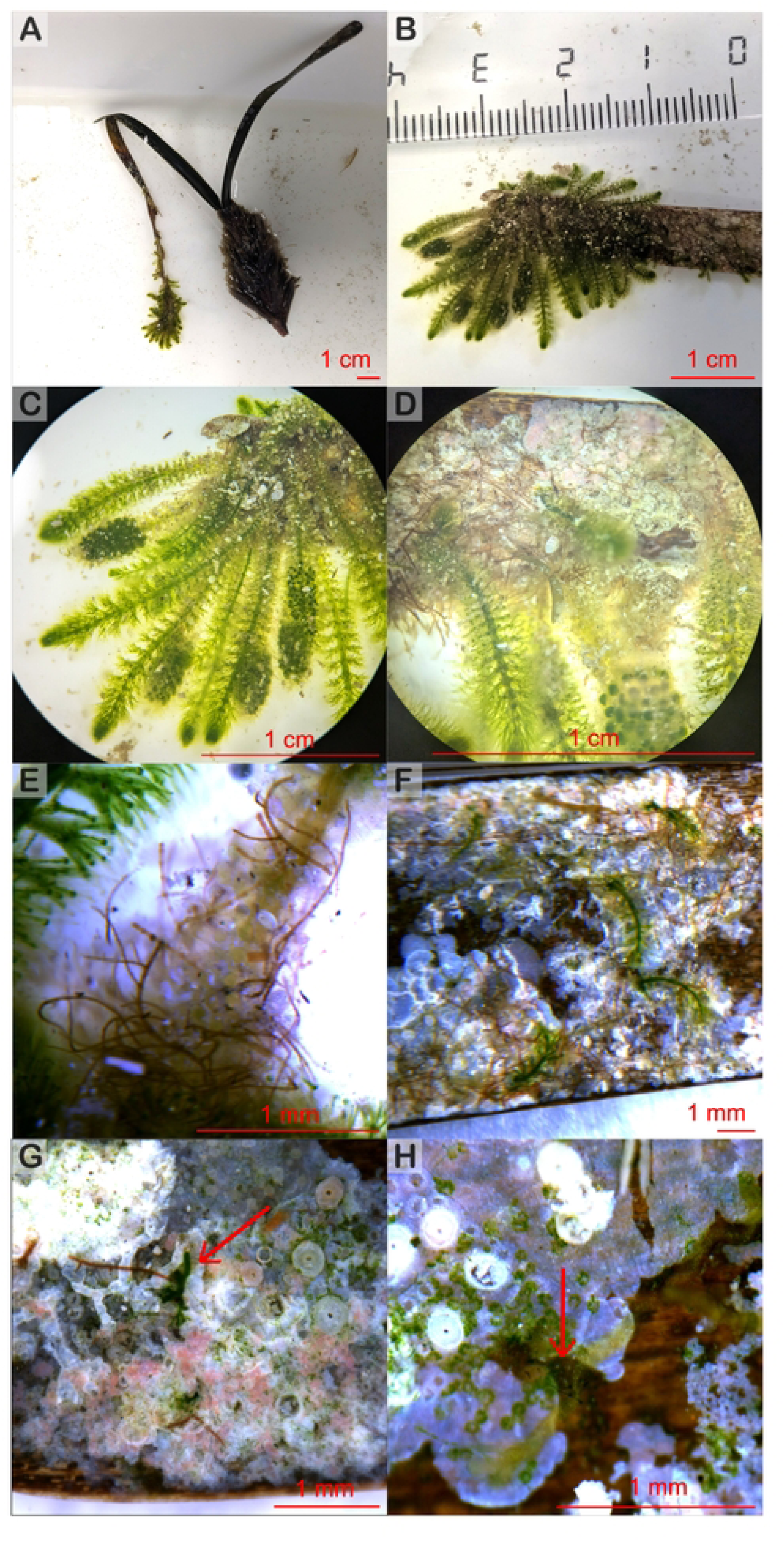
Progressive close-up of *B. occidentalis* development and attachment on *Posidonia oceanica* leaves. **A–D**) *B. occidentalis* consists of a bundle of green stalks and a brown rhizoid system that anchors onto the epiphytes present on *P. oceanica* leaves, primarily attaching near the top, where most of its biomass is concentrated. Stalks can be distinguished in vegetative ones (without gametophores) and fertile ones (with spherical dark-green gametophores). **E**) Sedimentation between the stalks occurs due to their fuzzy nature. **F-H**) Lower along the leaf fewer and smaller stalks are present, but an extensive rhizoid system may help the formation of additional stalks. The formation of new rhizoid and stalks - indicated by red arrows - starts from spot-like green patches. Photo credit: Silvia Paoletti.

## DISCUSSION

We detected a widespread presence of *B. occidentalis* in the Estany des Peix lagoon in 2023 and 2024, occupying all studied locations and habitats, including *Posidonia oceanica* seagrass meadows. Notably, we report for the first time *B. occidentalis* growing epiphytically on the leaves and rhizomes of *P. oceanica* (Figure 2E and 2F), reaching nearly 20% benthic coverage of this endemic habitat. The epiphytic nature of this exotic macroalgae could have detrimental effects on endemic *P. oceanica*. Most exotic species of macroalgae in the Mediterranean do not grow epiphytically on the blades of native macrophytes (e.g., seagrasses and macroalgae) but instead spread over sediment or grow over rhizomes in sparse meadows, like *Caulerpa racemosa* var. *cylindracea* on *P. oceanica* meadows (Ceccherelli et al. 2000; Ballesteros et al. 2007). An exception is the exotic rhodophyte *Lophocladia lallemandii*, which grows epiphytically on the *P. oceanica* leaves and has been shown to significantly increase seagrass mortality by 2.5–5 times compared to sites free of the invasive species (Marbà et al. 2014).

In Formentera, the exotic *B. occidentalis* grows abundantly on *Posidonia oceanica* leaves, covering nearly 25% of the leaf surface at all canopy levels—from the rhizomes to the upper parts of the seagrass (Figure 2E, 2F, and Figure 6). This extensive coverage could significantly reduce irradiance throughout the canopy. A similar negative interaction has been observed with *L. lallemandii*, which attenuates light within the *P. oceanica* canopy, limiting seagrass growth and decreasing shoot size and internode length (Marbà et al. 2014). Additionally, the epiphytic growth of *B. occidentalis* could damage *P. oceanica* leaves by increasing the weight of the epiphytic load. Seagrass leaves with heavy epiphyte cover have been shown to become more brittle and break off when epiphytic loads are high (0.8-1.5 g g^−1^ DW) or very high (>1.5 g g^−1^ DW) (Nelson 2017). In Estany des Peix, the epiphytic load of *B. occidentalis* on *P. oceanica* leaves ranged from 0 to 5.2 g g^−1^ leaf^−1^ DW, with almost 30 % (27 %) of leaves surpassing the 0.8 g g^−1^ DW threshold and 9 % supporting loads larger than 1.5 g g^−1^ DW of *B. occidentalis*. Field observations revealed that many *P. oceanica* leaves heavily overgrown by *B. occidentalis* (Figure 4A and Figure 6A) were bent, folded or partially torn, likely due to the excessive weight. Moreover, many detached *P. oceanica* leaves were found at the bottom of the lagoon, partially or fully covered with *B. occidentalis* (Figure 2B and 2H). Whether much of this epiphytic growth of *B. occidentalis* on the detrital leaves of *P. oceanica* preceded or followed detachment from the seagrass shoots remains to be determined.

*Batophora occidentalis* growing densely on the rhizomes and matte in sparse areas of *P. oceanica* with a 16.3 % cover in 2024 (Figure 2F) raises concerns for the native seagrass species. For example, the invasive *L. lallemandii* also colonizes rhizomes and leaves at the edges of meadows and in low-density *P. oceanica* patches, leading to reduced seagrass shoot size, leaf biomass, and a lower percentage of living shoots (Ballesteros et al. 2007). Similarly, sparse seagrass beds and meadow edges are negatively affected by the invasive green macroalga *C. cylindracea* (Ceccherelli et al. 2000). In the case of these two invasive species, their ability to attach to and overgrow the rhizomes and leaves of *P. oceanica* seems crucial to the invasion process (Ceccherelli et al. 2000; Ballesteros et al. 2007). Like invasive *C. cylindracea* and *Caulerpa taxifolia*, which use rhizoids to attach to the seagrass (Ballesteros et al. 2007), *B. occidentalis* has a holdfast with rhizoids as a means of attachment to structures (Terradas-Fernández et al. 2022). Potentially facilitated by the presence of other epiphytic organisms on *P. oceanica* leaves, this attachment mechanism likely allows *B. occidentalis* to start growing at the tip of leaves and spread downwards (Figure 6).

Quite alarmingly, we found a healthy stalk of *B. occidentalis* growing on a leaf of *P. oceanica* at the location outside the lagoon (Figure S4). Considering the sampling effort of this study, we calculated a potential density of 11.69 shoots m^-2^ of *P. oceanica* colonized by *B. occidentalis* in the meadow outside the lagoon. In 2023, *B. occidentalis* was also found inside three small rock pools at the entrance of the lagoon. Although in its native range the genus *Batophora* is found in a variety of habitats thanks to its eurythermal and euryhaline nature (Morrison 1984), distribution appears predominant in protected, unexposed areas, potentially in relation to wave energy (Nero and Sealey 2006). This was the case for the first recording of *B. occidentalis* in Formentera (Ballesteros 2020), however with time *B. occidentalis* may be successful in spreading to other suitable areas beyond the lagoon, posing a significant threat to the *P. oceanica* meadows of the Pityusic Islands, which are among the most extensive and well-preserved in the Mediterranean and a designated UNESCO World Heritage Site (Arnaud-Haond et al. 2012; del Valle Villalonga et al. 2023). The potential presence of *B. occidentalis* in other suitable areas around the Pityusic Islands is yet undetermined, nevertheless fundamental for the early detection of an ongoing expansion.

The Estany of Peix in Formentera is a sandy bottom lagoon with areas in the north-western part accumulating detrital leaves of *P. oceanica* in vast patches, meadows of *C. prolifera* and *C. nodosa* distributed throughout the lagoon and a meadow of *P. oceanica* at the entrance on the lagoon (Dantart et al. 1990). In the southern part of the lagoon (location 5 and 14; Figure 1A and 1B), *B. occidentalis* was growing on the sediment of *C. nodosa* meadows, where it became the dominant macrophyte in terms of both percentage cover and biomass (Figure S2, Figure 5A and 5B, Figure 6A and 6B). There is prior evidence of negative ecological effects caused by exotic macrophytes on the canopies of *C. nodosa*. For instance, in Sicily, the exotic seagrass *Halophila stipulacea* has been shown to reduce the shoot density of native *C. nodosa* year-round by forming a dense rhizome mat that outcompetes the native seagrass rhizomes, putatively pushing them down to the anoxic layer (Mannino et al. 2023; Mancuso et al. 2023). Similarly, in the Gulf of Naples, the invasive *C. cylindracea* has been found to reduce the photosynthetic performance of *C. nodosa* due to the phytotoxic effects of the secondary metabolite caulerpenyne produced by the invasive macroalga (Raniello et al. 2007). We also observed some evidence, albeit limited, of *B. occidentalis* growing on leaves of *C. nodosa* (Table 1 and Figure S5). The leaves of *C. nodosa* are smaller than those of *P. oceanica* (leaf surface of 9 and 83 cm^2^, respectively; (Duarte 1991)) and are likely unable to withstand heavy *B. occidentalis* loads, which may result in higher rates of leaf detachment or breakage compared to *P. oceanica*. Additionally, *C. nodosa* has a shorter average leaf lifespan of 55 days (Perez et al. 1994; Hemminga and Duarte 2000) compared to *P. oceanica,* which ranges from 202 - 345 days (Hemminga and Duarte 2000). Whether this 55-day period is sufficient for *B. occidentalis* to establish and grow on leaves of *C. nodosa* remains to be determined. In contrast, in the Mar Menor lagoon in Murcia, Terradas-Fernández et al. (2022) found no presence of *B. occidentalis* in *C. nodosa* meadows, suggesting that *B. occidentalis* may face challenges growing in dense seagrass canopies. We observed the opposite in Estany des Peix lagoon in Formentera, where *B. occidentalis* grew in *C. nodosa* meadows with a mean cover twice of that of *C. nodosa* (30 % vs 15.3 %, respectively in 2024; Table 1).

Similarly, *B. occidentalis* dominated both in percentage cover and biomass within *C. prolifera* meadows in the eastern part of the lagoon (location 7 and 13; Figure 1A and 1B, Figure S2, Figure 5A and 5B, Figure 6A and 6B). In two occasions *B. occidentalis* was found growing on the rhizome and leaves of *C. prolifera*, although in both instances the latter did not appear in healthy condition (Figure S4). The constraint on *B. occidentalis* in colonizing and covering *C. prolifera* may stem from several factors. *C. prolifera* is known for its rapid growth and regenerative capabilities, allowing it to outcompete native seagrasses for space in lagoonal environments (Pérez-Ruzafa et al. 2012; Parreira et al. 2021). Additionally, *Caulerpa* species produce caulerpenyne, a metabolite that serves as a chemical defense against herbivory and damage (Raniello et al. 2007), further enhancing their competitive advantage for space. This compound, combined with fast growth and a short blade turnover, may limit the overgrowth of *C. prolifera* by exotic *B. occidentalis*.

Our study represents the first record of quantitative benthic coverage and abundance of *B. occidentalis*. We document that in all surveyed habitats in the lagoon with the exception of *P. oceanica* meadows, *B. occidentalis* became the most abundant macrophyte in terms of benthic coverage and biomass: *Batophora occidentalis* cover was twice of that of *C. nodosa* (30 % vs 15.3 %, respectively in 2024), almost twice of that of *C. prolifera* (25.7 % vs 18 %, respectively in 2024) and the main macrophyte growing on sand (15.8 % in 2023). At times, the spread of exotic benthic algae can lead to complete substrate coverage, altering biodiversity, disrupting native assemblages and trophic interactions, and often resulting in habitat homogenization—ultimately leading to the overall degradation of littoral ecosystems (Piazzi and Balata, 2009; Borriglione et al. 2024). As an example, *Caulerpa cylindracea* and *Womersleyella setacea* invaded a range of Mediterranean habitats becoming the main macrophyte species causing varying degrees – but always negative – of habitat impoverishment (Piazzi and Balata, 2009). To extreme extents this happened with the brown algae *Rugulopteryx okamurae*, which exceeded 85 % coverage in certain areas of the Strait of Gibraltar (García-Gómez et al. 2021) and reached 100% coverage in areas of the North-western Mediterranean (Borriglione et al. 2024). The high coverage values of *B. occidentalis* observed at Estany des Peix are concerning, as it may gradually overtake benthic substrates over time.

Exotic *B. occidentalis* was also found growing on several natural hard substrates such as rocks, pebbles, and dead shells of gastropods and bivalves (Figure 2). Similarly, Forteza et al. (2024) documented *B. occidentalis* growing on all sediment types from muddy sand to rocks along Estany of Peix perimeter, while Terradas-Fernández et al. (2022) documented *B. occidentalis* growing throughout the north-eastern perimeter of Mar Menor, attached to several substrates including pebbles and mollusks like *Hexaples trunculus* and *Pinna nobilis*. Recently in the Chesapeake Bay (USA), introduced *Batophora oerstedii* was found growing attached to shells in shallow water less than 20 cm depth (Hall and Schneider 2023). However, it remains undocumented whether *B. occidentalis* can grow on living hard-shell species such as gastropods or bivalves. If it does, this could have negative implications for these animals, as seen in bivalves affected by epibiont fouling (Garner and Litvaitis, 2016). We also found *B. occidentalis* in the Estany des Peix growing on artificial hard substrates such as wooden peers (Figure 3), as documented in the Chesapeake Bay (Hall and Schneider 2023), plastic debris, as documented in Mar Menor (Terradas-Fernández et al. 2022), and metallic substrates, including can debris, boat chains and boat hulls (Figure 3), as reported by Forteza et al. (2024). Specifically, we found *B. occidentalis* growing in almost 20 % (17.4 %) of the hulls of the boats anchored to the main marina of the lagoon in 2024. This is worrisome, as these small boats could act as vectors to spread *B. occidentalis* to other marinas and locations in the Mediterranean. Vessel hull fouling is a primary vector by which invasive species are transported globally (Costello et al. 2022). A recent study by Ulman et al. (2019) on the spread of exotic species via recreational boats in the Mediterranean Sea found that 71% of sampled vessels hosted at least one, and up to 11, exotic species on their hulls and the majority of these vessels visited distant marinas where such exotic species were not yet present, highlighting their role as vectors for biological invasions.

Some species of exotic macroalgae are easily identified by their morphological attributes. For instance, exotic *Codium fragile* can be distinguished from native *Codium decorticatum* by the presence of apiculate utricle tips visible in cross sections (Geraldi et al. 2014). In the case of *B. occidentalis*, the species can be distinguished from the native relative *Dasycladus vermicularis* by the distance between whorls and the distribution of gametophores in the whorl branchlets (Terradas-Fernández et al. 2022), but not the same applies to the different species within the genus. *Batophora occidentalis* populations in Formentera and Mar Menor have been suggested to belong to the same species - and possibly same introduction event - due to overlapping morphological traits (Terradas-Fernández et al. 2022). The specimens have been tentatively identified as *Batophora occidentalis* or *Batophora occidentalis* var. *largoensis* in the Mediterranean through morphological literature review (Ballesteros et al. 2020; Terradas-Fernández et al. (2022), however the assignment of a specific species or variety was not possible due to the current scarcity of molecular data available on this genus. Further genetic investigation is needed to identify the specimens as a key aspect to understand the species ecology.

Primary producers are ranked as the most damaging group of marine exotics based on their quantified ecological impacts, often exerting negative effects on other primary producers (Anton et al. 2019, 2020a). In this study, we document *B. occidentalis* growing on the leaves and rhizomes of the endangered seagrass *P. oceanica*, as well as on the meadows of *C. nodosa* and *C. prolifera*. The most harmful effects of exotic species become apparent when they replace ecosystem engineers (Crooks 2002). The endemic seagrass *P. oceanica* is a key engineering species in the Mediterranean Sea (Green and Short 2003), covering an estimated area of 1.2 million hectares (Telesca et al., 2015). These meadows provide critical habitat for marine biodiversity, stabilize sediments, and contribute to carbon sequestration. These functions have led to the designation of *P. oceanica* meadows as habitats of priority interest under the European Habitats Directive (92/43/CEE). Similarly, meadows of *C. nodosa* provide relevant ecosystem services acting as nursery and feeding grounds, reducing coastal erosion and supporting nutrient cycling and carbon sequestration (Da Ros et al. 2021). Finally, the meadows of *C. prolifera* have a key role in sinking dissolved inorganic nitrogen helping resistance mechanisms to eutrophication in temperate coastal lagoons (Bernardeau-Esteller et al. 2023). In the present study, we provide evidence of the new potential ecological threats posed by *B. occidentalis* to these important native biogenic habitats and we provide recommendations for early management.

Four years after its first detection in the Estany des Peix, *B. occidentalis* is now widely distributed across all benthic habitats within the lagoon, with observations suggesting a potential spread beyond its boundaries. While eradication is often proposed for localized invasions (Thresher & Kuris, 2004), success stories in marine ecosystems remain scarce (e.g., *Spartina alterniflora*; Kerr, 2024). Although the exotic *B. occidentalis* in Formentera is currently confined to the small lagoon (approximately 2 km²), it has achieved extensive coverage, with an average density of 5,400 stalks per square meter. As a result, total eradication may not be feasible without significant effort and financial investment. However, targeted removal efforts could be performed focusing on the lagoon’s entrance to prevent further natural spread. To mitigate the risk of *B. occidentalis* spreading to distant locations, likely transported by boats permanently or temporarily moored in the lagoon, prevention strategies should be implemented, including periodic mandatory cleaning of biofouling from boat hulls in Estany des Peix. Additionally, key ecological information necessary for effective management, such as its reproductive cycle and growth seasonal patterns, remains unknown and requires further investigation. In this paper, we advocate for two urgent actions. First, we call for intensified research on *B. occidentalis*, including monitoring lagoonal environments and marinas across the Balearic Islands for early detection of spread. Second, we urge local and regional governments to take immediate management action by prioritizing the removal of *B. occidentalis* from boat hulls within the lagoon and clearing it near the lagoon’s inlet to prevent further colonization in Formentera and other Mediterranean locations.

## Acknowledgements

Anton A. was supported by Ramon y Cajal grant (number RYC2021-033047-I), funded by MCIN/AEI/10.13039/501100011033 and by the European Union ‘NextGenerationEU/PRTR,’ by the Universitat de les Illes Balears, and by the ‘Pla Anual d’Impuls del Turisme Sostenible per al període 2023’ of the Balearic Government. This research was carried out within the framework of the “Maria de Maeztu Excellence Unit” accreditation of IMEDEA, Grant CEX2021-001198 funded by MCIN/AEI/10.13039/501100011033. Paoletti S received the support of an INPHINIT fellowship from the “la Caixa” Foundation (ID 100010434). The fellowship code is “LCF/BQ/DI24/12070025”. We thank Balearia for sponsoring the ferry trips from Mallorca to Formentera. The Spanish Ministry of Science, the Universitat de les Illes Balears, and Balearia accept no responsibility for the opinions, statements and contents included in the project and/or the results thereof, which are entirely the responsibility of the authors.

## Data availability

The dataset related to this article is available in Zenodo for peer-review purpose here: https://zenodo.org/records/15130273?preview=1&token=eyJhbGciOiJIUzUxMiIsImlhdCI6MTc0MzY3NTg4OSwiZXhwIjoxNzU2NTExOTk5fQ.eyJpZCI6IjNmNTcxNmZmLWJkZWYtNGY4MS1iMGU0LTNlNmZlNzY0ZWU0YSIsImRhdGEiOnt9LCJyYW5kb20iOiIwNzNjOTI0ZjRmMTY0MmFmMGVlMGI1YjVjNjdiMjBhZiJ9.lmYyKHuXbPlxtvk4vom7S-h47JEEh6mFwpBut_GlJVW589dFQFKnO3DgYdzN5aN7ff9cqFvXQteg2Xd4RIhGyg

Once the article is accepted for publication, the dataset will be freely accessible as open access.

## Notes

### Competing Interest Statement

The authors have declared no competing interest.

